# A practical generation interval-based approach to inferring the strength of epidemics from their speed

**DOI:** 10.1101/312397

**Authors:** Sang Woo Park, David Champredon, Joshua S. Weitz, Jonathan Dushoff

## Abstract

Infectious-disease outbreaks are often characterized by the reproductive number and 𝓡 exponential rate of growth *r*. 𝓡 provides information about outbreak control and predicted final size. Directly estimating 𝓡 is difficult, while *r* can often be estimated from incidence data. These quantities are linked by the generation interval – the time between when an individual is infected by an infector, and when that infector was infected. It is often infeasible to obtain the exact shape of a generation-interval distribution, and to understand how this shape affects estimates of 𝓡. We show that estimating generation interval mean and variance provides insight into the relationship between and 𝓡 *r*. We use examples based on Ebola, rabies and measles to explore approximations based on gamma-distributed generation intervals, and find that use of these simple approximations are often sufficient to capture the *r*–*𝓡* relationship and provide robust estimates of *𝓡*.

## 1. Introduction

Infectious disease research often focuses on estimating the reproductive number, i.e., the number of new infections caused on average by a single infection. This number is termed the reproductive number – 𝓡. The reproductive number provides information about the disease’s potential for spread and the difficulty of control. It is described in terms of an average [4] or an appropriate sort of weighted average [9].

The reproductive number has remained a focal point for research because it provides information about how a disease spreads in a population, on the scale of disease generations. As it is a unitless quantity, it does not, however, contain information about *time*. Hence, another important quantity is the population-level *rate of spread, r*. The initial rate of spread can often be measured robustly early in an epidemic, since the number of incident cases at time *t* is expected to follow *i*(*t*) *≈ i*(0) exp(*rt*). The rate of growth can also be described using the “characteristic time” of exponential growth *C* = 1*/r*. This is closely related to, and simpler mathematically than, the more commonly used doubling time (given by *T*_2_ = ln(2)*C ≈* 0.69*C*).

In disease outbreaks, the rate of spread, *r*, can be inferred from case-incidence reports, e.g., by fitting an exponential function to the incidence curve [27, 29, 24]. Estimates of the initial exponential rate of spread, *r*, can then be combined with a mechanistic model that includes unobserved features of the disease to esimate the initial reproductive number, 𝓡. In particular, 𝓡 can be calculated from *r* and the generation-interval distribution using the generating function approach popularized by [44].

The *generation interval* is the amount of time between when an individual is infected by an infector, and the time that the infector was infected [39]. While *r* measures the speed of the disease at the population level, the generation interval measures speed at the individual level. Generation interval distributions are typically inferred from contact tracing, sometimes in combination with clinical data [7, 20, 16]. Generation interval distributions can be difficult to ascertain empirically [29, 8], and the generation-function approach depends on an entire distribution. Multiple studies have explored how general-interval distributions affect the *r*– 𝓡 relationship (summarized in Table 1). However, the use of an entire distribution makes it difficult to determine which features of the distributions are essential to connect measurements of the rate of spread *r*, with the reproductive number, 𝓡. Moreover, different assumptions about the shape of generation interval distributions have led to seemingly contradictory results [46, 37].

**Table 1:** Summary of previous studies that relate growth rate, *r*, with reproductive number, 𝓡, using generation interval distributions.

| Method | Required information | Assumptions/Notes | References |
| --- | --- | --- | --- |
| Delta approximation | • Mean generation interval | • Assumes fixed generation interval<br>• Provides an upper bound for $\mathcal{R}$ estimate | [45, 37, 26, 42] |
| Empirical (histogram-based) | • Generation interval samples | • Relies on binning generation intervals into discrete histograms | [45] |
| Empirical (sample-based) | • Generation interval samples | • Uses discretized Euler-Lotka equation | [14] |
| Euler-Lotka | • Entire generation interval distribution | • Uses moment generating function | [23] |
| Exponential approximation (SIR) | • Mean generation interval | • Assumes exponentially distributed generation interval<br>• Predicts linear relationship between $r$ and $\mathcal{R}$ | [6, 33, 12, 45, 42] |
| Gamma approximation | • Mean/SD generation interval | • Assumes gamma distributed generation intervals | [29, 25, 42, 31] |
| Normal approximation | • Mean/SD generation interval | • Assumes normally distributed generation intervals<br>• Predicts decreasing $r$-$\mathcal{R}$ relationship for high $r$ | [45, 42] |
| SEIR (gamma) | • Mean/SD latent period | • Assumes gamma distributed latent and infectious periods<br>• See [48] for summary of all special cases | [3, 46, 45, 37] |
| SEIR (generalized) | • Laplace transformation of latent period distribution<br>• Laplace transformation of infectious period distribution | • Independent latent and infectious periods<br>• Generalizes gamma distributed latent and infectious periods | [48] |
| Trapezoid approximation | • Period of no infection after exposure<br>• Time until maximum infectivity<br>• Duration of maximum infectivity<br>• Time until recovery | • Assumes trapezoid shaped infection kernel | [37] |

The primary goal of this paper is to further explore and explain the *r*– 𝓡 relationship. We explore the qualitative relationship between generation time, initial rate of spread *r*, and initial reproductive number 𝓡 using means, variance measures and gamma approximations. We show that summarizing generation-interval distributions using moments provides biological explanations that unify previous findings. We further discuss the generality of the gamma approximation and provide examples of its relevance in using realistic epidemiological parameters from previous outbreaks. By doing so, we shed light on the underpinnings of the relationship between *r* and 𝓡, and on the factors underlying its robustness and its practical use when data on generation intervals is limited or hard to obtain.

## 2. Relating 𝓡 and *r*

In this section, we introduce an analytical framewok, and recapitulate previous work relating growth rate *r* and reproductive number 𝓡. These two quantities are linked by the generation-interval distribution, which describes the interval between the time an individual becomes *infected* and the time that they *infect* another individual. In particular, if 𝓡 is known, a shorter generation interval means a faster epidemic (larger *r*). Conversely (and perhaps counter-intuitively), if *r* is known, then shorter disease generations imply a *lower* value of 𝓡, because more *individual* generations are required to realize the same *population* spread of disease [11, 32] (see Fig. 1).

**Figure 1:**
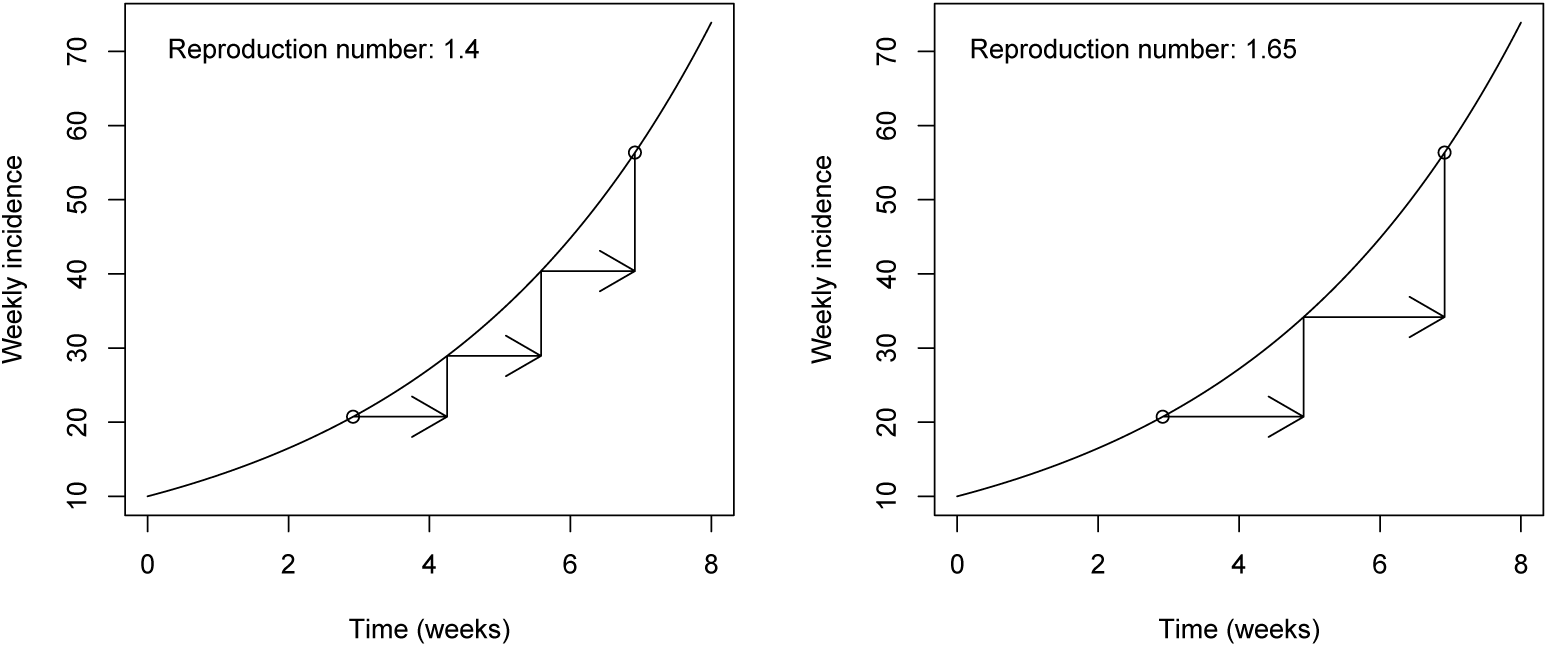
Two hypothetical epidemics with the same growth rate (*r* = 0.25*/* week) and fixed generation intervals. Assuming a short generation interval (fast transmission at the individual level) implies a smaller reproductive number 𝓡_0_ (panel A) when compared to a longer generation interval (slow transmission at the individual level, panel B).

The generation-interval distribution can be defined using a renewal-equation approach. A wide range of disease models can be described using the model [15, 10, 36, 1, 44, 37]:

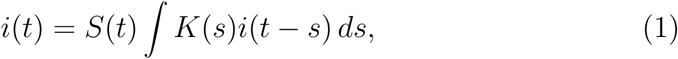

where *t* is time, *i*(*t*) is the incidence of new infections, *S*(*t*) is the *proportion* of the population susceptible, and *K*(*s*) is the intrinsic infectiousness of individuals who have been infected for a length of time *s*.

The *basic* reproductive number, defined as the average number of secondary cases generated by a primary case in a fully susceptible population [4, 9], is

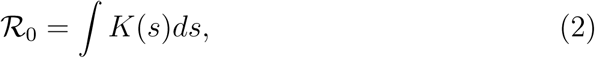

and the *intrinsic* generation-interval distribution is

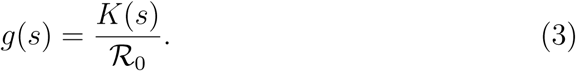

The “intrinsic” interval can be distinguished from “realized” intervals, which can look “forward” or “backward” in time [8] (see also earlier work [39, 28]). In particular, it is important to correct for biases that shorten the intrinsic interval when generation intervals are observed through contact tracing during an outbreak.

In this model, disease growth is predicted to be approximately exponential in the early phase of an epidemic, because the depletion in the effective number of susceptibles is relatively small. Thus, for the exponential phase, we write:

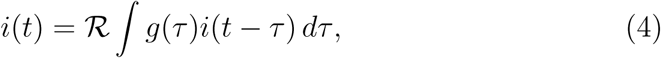

where 𝓡 = 𝓡 _0_*S* is the *effective* reproductive number. We can then solve for the characteristic time *C* by assuming that the population is growing exponentially: i.e., substitute *i*(*t*) = *i*(0) exp(*t/C*) to obtain the exact speed-strength relationship [10]:

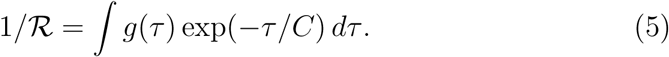

This fundamental relationship dates back to the work of Euler and Lotka [23]. We will explore the shape of this relationship using parameters based on human infectious diseases, and investigate approximations based on gamma-distributed generation intervals.

## 3. Approximation framework

### 3.1. Approximation method, in theory

We do not expect to know the full distribution *g*(*τ*) – particularly while an epidemic is ongoing – so we are interested in approximations to 𝓡 based on limited information. We follow the approach of [29] and approximate the generation interval with a gamma distribution. We prefer the gamma distribution to the standard normal approximation used in many applications for a number of reasons. First, it is more biologically realistic, since it is confined to non-negative values. Second, it has a convenient moment-generating function, and provides a corresponding simple form for the *r*– 𝓡 relationship that can be parameterized with only two parameters that have biologically relevant meanings that can assist in explaining intuition behind the *r*– 𝓡 relationship. Third, it generalizes the result obtained from simple Susceptible-Infectious-Recovered (SIR) models, and approximately matches Susceptible-Exposed-Infectious-Recovered (SEIR), in the case where the latent period and infectious period are similar (see Appendix). While the gamma approximation has been applied to infer 𝓡 in previous outbreaks (Table 1), its theoretical properties and practical importance has not yet been explored in depth.

For biological interpretability, we describe the gamma distribution using the mean 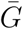 and the squared coefficient of variation *κ* (thus *κ* = 1*/a*, and 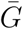, where *a* and *θ* are the shape and scale parameters under the standard parameterization of the gamma distribution). Substituting the gamma distribution into (5) then yields the gamma-approximated speed-strength re-lationship:

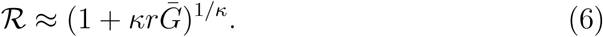

We write:

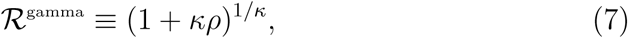

where 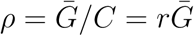 measures how fast the epidemic is growing (on the time scale of the mean generation interval) – or, equivalently, the length of the mean generation interval (in units of the characteristic time of exponential growth). The longer the generation interval is compared to *C*, the higher the estimate of 𝓡 (see Fig. 1). We then explore the behaviour of the gamma-approximated speed-strength relationship 𝓡 ^gamma^ defined above (equivalent to the Tsallis “q-exponential”, with *q* = 1 *-κ* [43]): its shape determines how the estimate of 𝓡 changes with the estimate of normalized generation length *ρ*.

For small *ρ*, 𝓡 ^gamma^ always looks like 1 + *ρ*, regardless of the shape parameter 1*/κ*, which determines the curvature: if 1*/κ* = 1, we get a straight line, for 1*/κ* = 2 the curve is quadratic, and so on (see Fig. 2). For large values of 1*/κ*, 𝓡 ^gamma^ converges toward exp(*ρ*).

**Figure 2:**
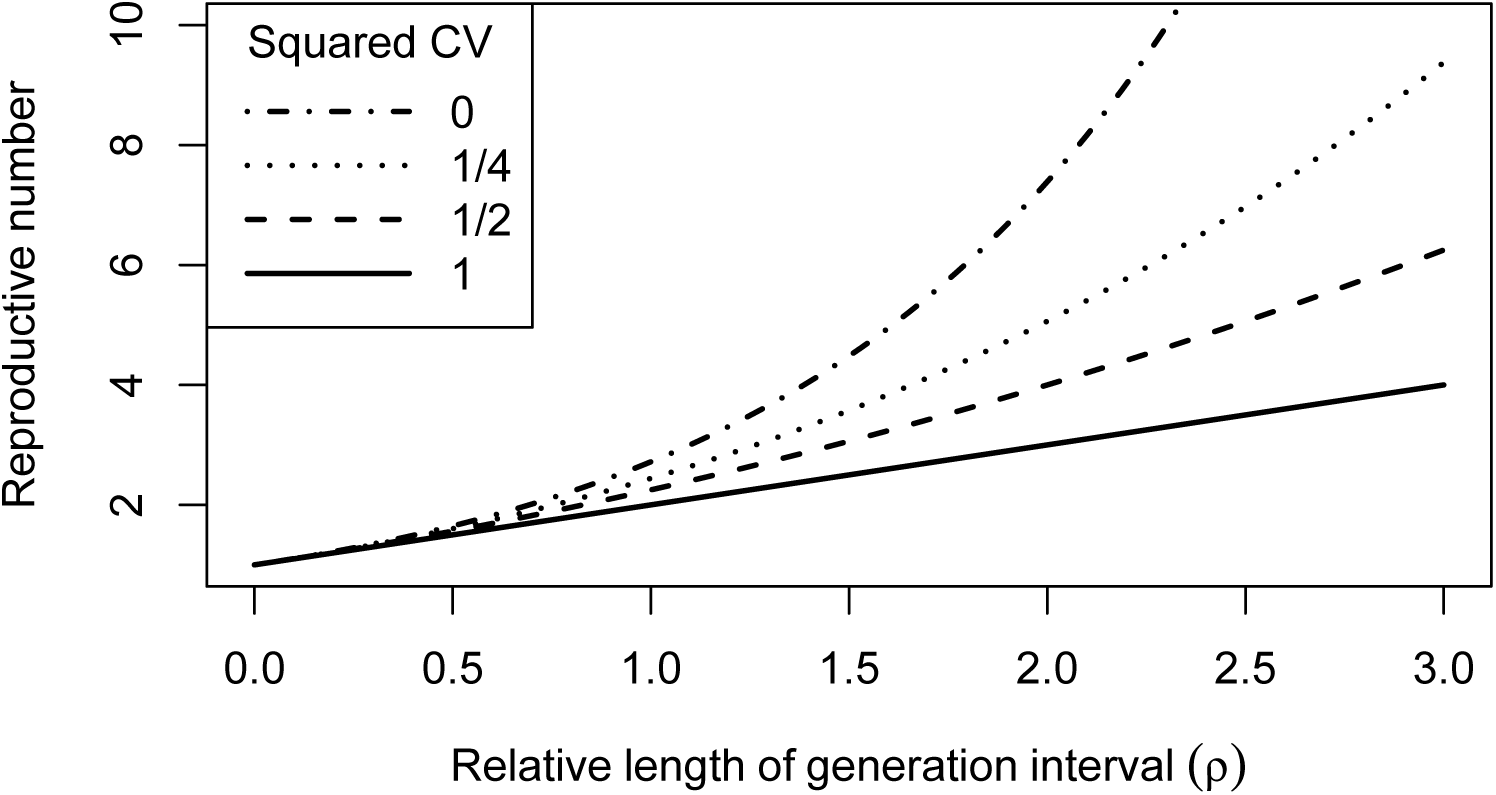
The approximate relationship (6) between mean generation time (relative to the characteristic time of exponential growth, 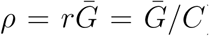 and reproductive number. The curves correspond to different amounts of variation in the generation-interval distribution.

The limit as *κ →* 0 is reasonably easy to interpret. The incidence is increasing by a factor of exp(*ρ*) in the time it takes for an average disease generation. If *κ* = 0, the generation interval is fixed, so the average case must cause exactly 𝓡 = exp(*ρ*) new cases. If variation in the generation time (i.e., *κ*) increases, then some new cases will be produced before, and some after, the mean generation time. Since we assume the disease is increasing exponentially, infections that occur early on represent a larger proportion of the population, and thus will have a disproportionate effect: individuals don’t have to produce as many lifetime infections to sustain the growth rate, and thus we expect 𝓡 *<* exp(*ρ*) (as shown by [44]).

The straight-line relationship for *κ* = 1 also has a biological interpretation. This case corresponds to the classic SIR model, where the infectious period is exponentially distributed [6]. In this case, recovery rate and infection rate are constant for each individual. The rate of exponential growth per generation is then given directly by the net per capita increase in infections: 𝓡 *-* 1, where one represents the recovery of the initial infectious individual.

Characterizing the *r*– 𝓡 relationship with mean and coefficient of variation also helps explain results based on compartmental models [46, 37], because the mean and variance of the generation interval is linked to the mean and variance of latent and infectious periods. Shorter mean latent and infectious periods result in shorter generation intervals, and thus smaller 𝓡 estimates. Less-variable latent periods result in less-variable generation intervals, and thus higher 𝓡 estimates. Less-variable infectious periods have more complicated effects. While they reduce variation in generation interval (which would be expected to increase 𝓡 estimates), they also reduce the mean generation interval [39] (which would lead to a decrease). This explains the apparent anomaly between earlier results: when mean generation interval is held constant, the effect of less-variable infectious periods is to reduce variation, leading to lower 𝓡 for a given *r* [37]; when mean infectious period is held constant the effect of reducing generation interval will be stronger, leading to higher 𝓡 [46].

There is a simple intuition for this latter result. In an exponentially growing epidemic, things that happen earlier have the largest effect. If we increase the variation in latent period while holding 𝓡 constant, we have more early progression and more late progression to infectiousness. The former effect will be more important, and thus increasing variation should increase *r* for a given value of 𝓡 – or, equivalently, decrease 𝓡 for a given value of *r*. Increasing variation in infectious period has the opposite effect: it increases the amount of both early and late recovery, and thus leads to smaller *r* for a given value of 𝓡 (and conversely).

### 3.2. Approximation method, in practice

We test our approximation method by generating a pseudo-realistic generation-interval distributions using previously estimated/observed latent and infectious period distributions for different diseases (Table 2). For each pseudo-realistic distribution, we calculate the “true” relationship between *r* and 𝓡 and compare it with a relationship inferred based on gamma distribution approximations. These approximations are first done with large amounts of data, allowing us to evaluate how well the approximations describe the *r*– 𝓡 relationship under ideal conditions, and then tested with smaller amounts of data to assess how well methods perform when data is limited.

**Table 2:** Parameters that were used to obtain theoretical generation distri-butions for each disease. Reproduction numbers are represented as points in figure Fig. 3–5. Ebola parameters in triples represent Sierra Leone, Liberia, Guinea.

| Disease | Ebola | Measles | Rabies |
| --- | --- | --- | --- |
| Parameter | Values |  |  |
| Reproduction number | 1.38, 1.51, 1.81 [7] | 12.5-18 [5] | 1.05-1.32 [13] |
| Mean incubation period (days) | 11.4 [7] | 12.77 [19] | 24.24 [13] |
| SD incubation period (days) | 8.1 (see Sec. 4.1) | 2.67 [19] | 29.49 [13] |
| Mean infectious period (days) | 5 [7] | 6.5 [5] | 3.57 [13] |
| SD infectious period (days) | 4.7 [7] | 2.9 (see Sec. 4.2) | 2.26 [13] |
| Relative length of generation interval ($\rho$) | 0.35, 0.45, 0.68 [7] | NA | NA |
| Mean generation interval (days) | 16.2 | 15.0 | 26.6 |
| CV generation interval | 0.58 | 0.21 | 1.09 |

Estimating generation intervals is complex; our goal with pseudo-realistic distributions is not to precisely match real diseases, but to generate distributions that are likely to be roughly as challenging for our approximation methods as real distributions would be. We construct pseudo-realistic intervals from sampled latent and infectious periods by adding the sampled latent period to a an infection delay chosen uniformly from the sampled infectious period:

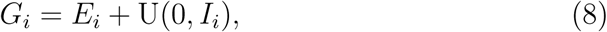

where *G*_*i*_, *E*_*i*_ and *I*_*i*_ are the sampled intrinsic generation interval, latent period, and infectious period, respectively, and U represents a uniform random deviate. This implicitly assumes that infectiousness is constant across the infectious period [13]. We sample from latent and infectious periods obtained from observations (for empirical distributions), or by using a uniform set of quantiles (for parametric distributions). For the purpose of constructing pseudo-realistic distributions, we do *not* attempt to correct for the fact that observed intervals may be sampled in a context more relevant to backward than to intrinsic generation intervals (see [8]). We sample latent periods at random, and infectious periods by length-weighted resampling (since longer infectious period implies more opportunities to infect). For our examples, we used 10,000 quantiles for each parametric distribution and 10,000 sampled generation intervals for each disease (see Appendix).

We then calculate “exact” relationships (for our pseudo-realistic distributions) by substituting sampled generation intervals into the exact speed-strength relationship (5). This relationship is then compared to the corresponding gamma-approximated relationship (6).

All calculations, numerical analyses and figures were made with the software platform R [34]. Code is freely available at https://github.com/dushoff/ link calculations.

## 4 Results

We investigate this approximation approach using three different examples. These examples also serve to demonstrate that robust estimates could be made with less data and potentially earlier in an outbreak – a point we revisit in the Discussion. Our initial investigation of this question was motivated by work on the West African Ebola Outbreak [47], so we start with that example. To probe the approximation more thoroughly, we also chose one disease with high variation in generation interval (canine rabies), and one with a high reproductive number (measles). For simplicity, we assumed that latent and infectious periods are equivalent to incubation and symptomatic periods for Ebola virus disease (EVD), measles, and canine rabies.

### 4.1 Ebola

We generated a pseudo-realistic generation-interval distribution for Ebola virus disease (EVD) using information from [7] and a lognormal assumption for both the incubation and infectious periods. In contrast to gamma distributed incubation and infectious periods assumed by [7], we used a lognormal assumption for our components because it is straightforward and should provide a challenging test of our gamma approximation (see Appendix for results using gamma components). We used the reported standard deviation for the infectious period, and chose the standard deviation for the incubation period to match the reported coefficient of variation for the serial interval distribution, since this value is available and is expected to be similar to the generation interval distribution for EVD [7].

We then used our pseudo-realistic distribution to calculate both the exact (5) and gamma-approximated (6) speed-strength relationships (see Fig. 3). The approximation is within 1% of the pseudo-realistic distribution it is approximating across the range of country estimates, and within 5% across the range shown. It is also within 2% of the World Health Organization (WHO) estimates derived from the observed time series assuming a poisson process with a known serial interval distribution.

**Figure 3:**
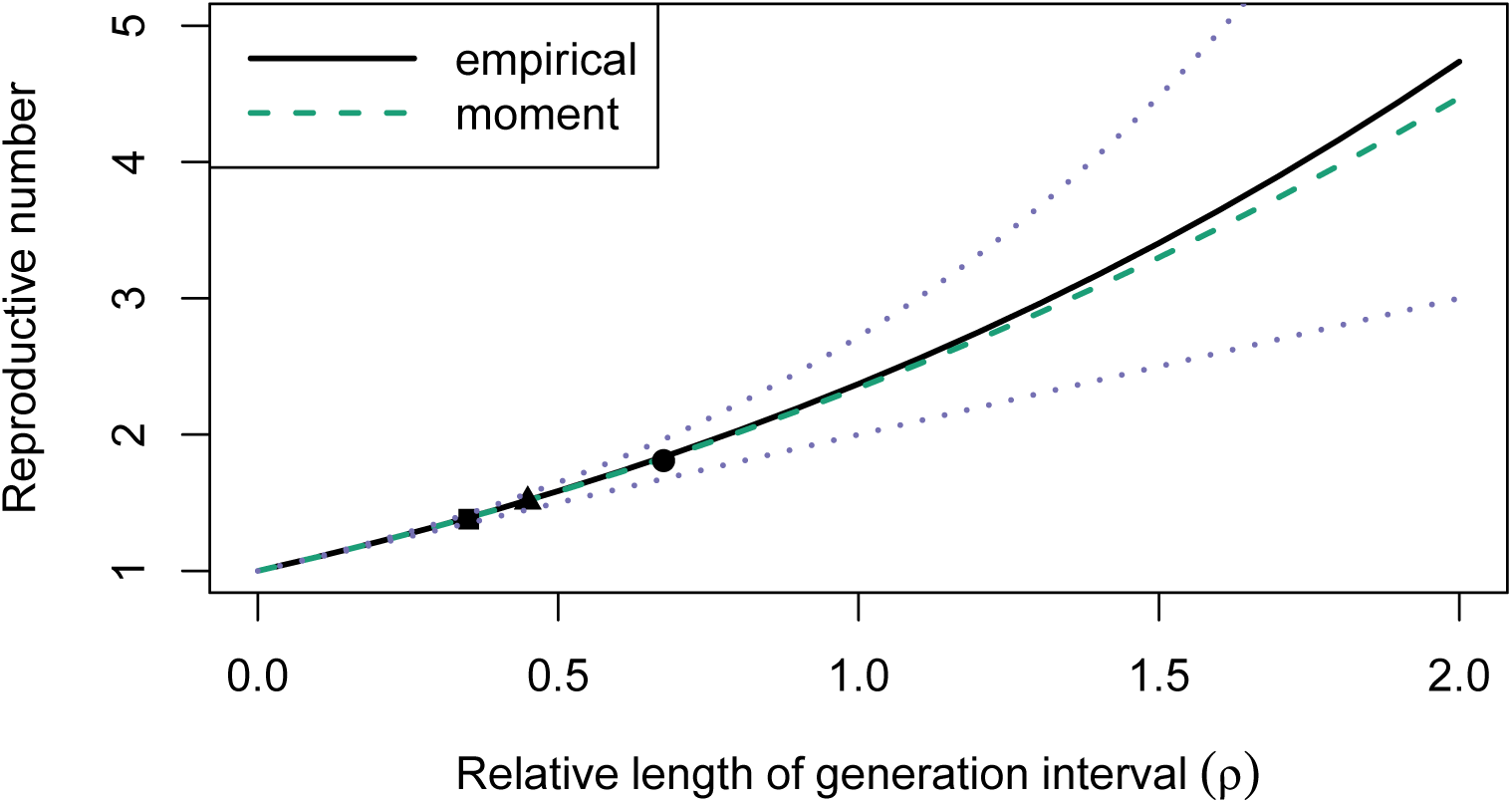
Estimating 𝓡 for the West African Ebola Outbreak. (solid curve) The exact speed-strength relationship (5) using a pseudo-realistic generation-interval distribution based on [7]. (dashed curve) The Gamma approximated speed-strength relationship (6), using the mean and CV of a pseudo-realistic generation-interval distribution. (dotted curves) Naive approximations based on exponential (lower) and fixed (upper) generation distributions. Points indicate estimates for the three focal countries of the West African Ebola Outbreak calculated by [7]: Sierra Leone (square, 𝓡 = 1.38), Liberia (triangle, 𝓡 = 1.51), and Guinea (circle, 𝓡 = 1.81). Initial growth rate for each outbreak was inferred from doubling periods reported by [7] (*r* = ln(2)*/T*_2_).

### 4.2 Measles

We also applied the moment approximation to a pseudo-realistic generation-interval distribution based on information about measles from [19], [22], and [5]. Incubation periods were assumed to follow a lognormal distribution [19]. Infectious periods were assumed to follow a gamma distribution with coefficient of variation of 0.2 [38, 22, 17]. Since variation in infectious period is relatively low [38, 17], and infectious period is short compared to incubation period, this choice is reasonable (and our results are not sensitive to the details).

Here, we found surprisingly close agreement between the exact and approximate relationships between *r* and 𝓡 across a much wider range of interest (a difference of *<* 1% for 𝓡 up to *>* 20) (see Fig. 4). On examination, this closer agreement is due to the smaller overall variation in generation times in measles: when overall variation is small, differences between distributions have less effect.

**Figure 4:**
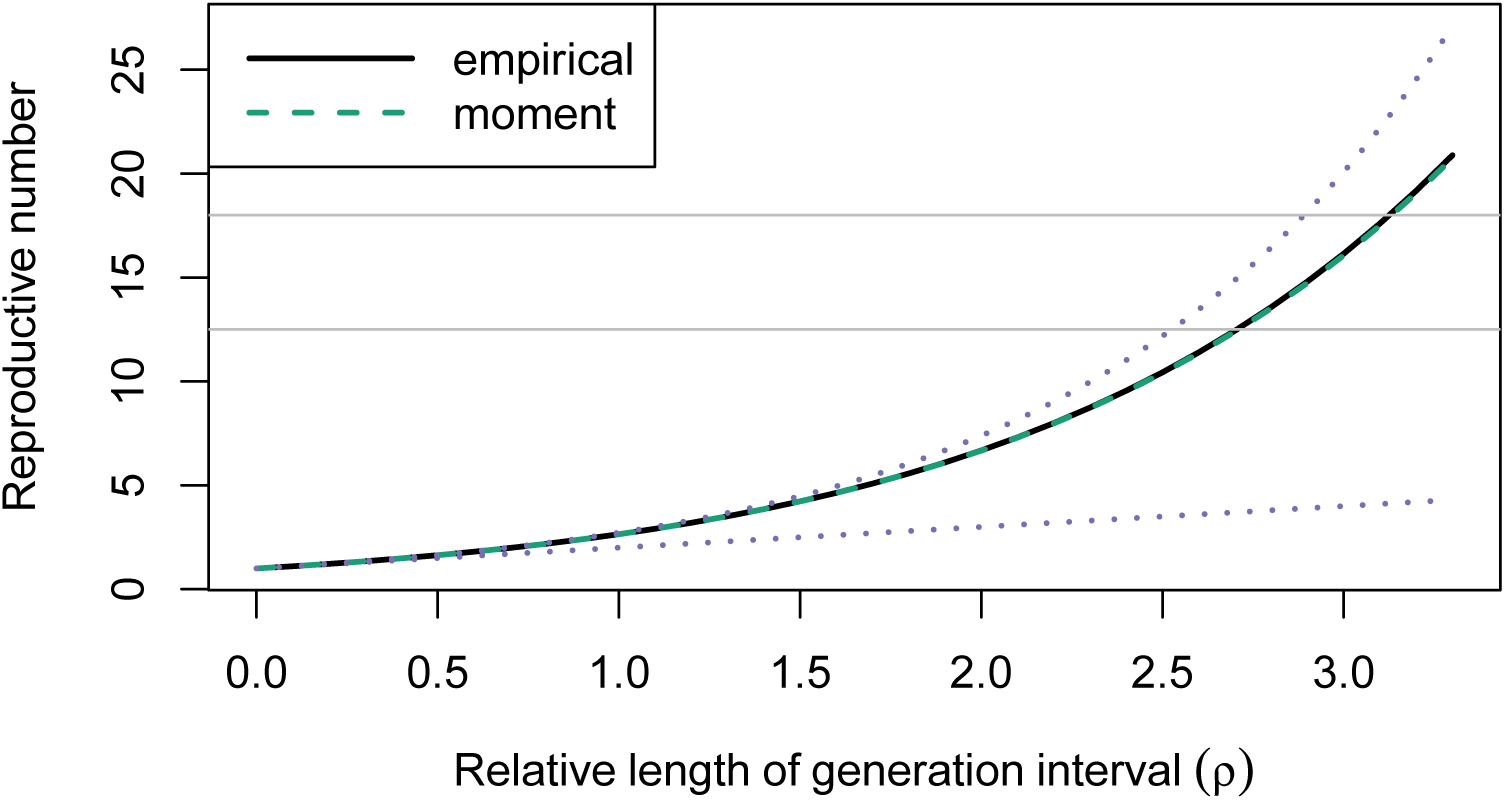
Estimating 𝓡 for measles. (solid curve) The exact speed-strength relationship (5) using a pseudo-realistic generation-interval distribution. (dashed curve) The gamma-approximated relationship (6) using the mean and CV of a pseudo-realistic generation-interval distribution (this curve is almost invisible because it overlaps the solid curve) (dotted curves) Naive approximations based on exponential (lower) and fixed (upper) generation distributions. Gray horizontal lines represent 𝓡 ranges estimated by [5]: 12.5-18.

### 4.3 Rabies

We did a similar analysis for rabies by constructing a pseudo-realistic generation-interval distribution from observed incubation and infectious period distributions (see Fig. 5). Since estimates of 𝓡 for rabies are near 1, there is small difference between the naive estimates and the gamma approximated speed-strength relationship. But, looking at the relationship more broadly, we see that the moment-based approximation would do a poor job of predicting the relationship for intermediate or large values of 𝓡 – in fact, a poorer job than if we use the approximation based on exponentially distributed generation times.

**Figure 5:**
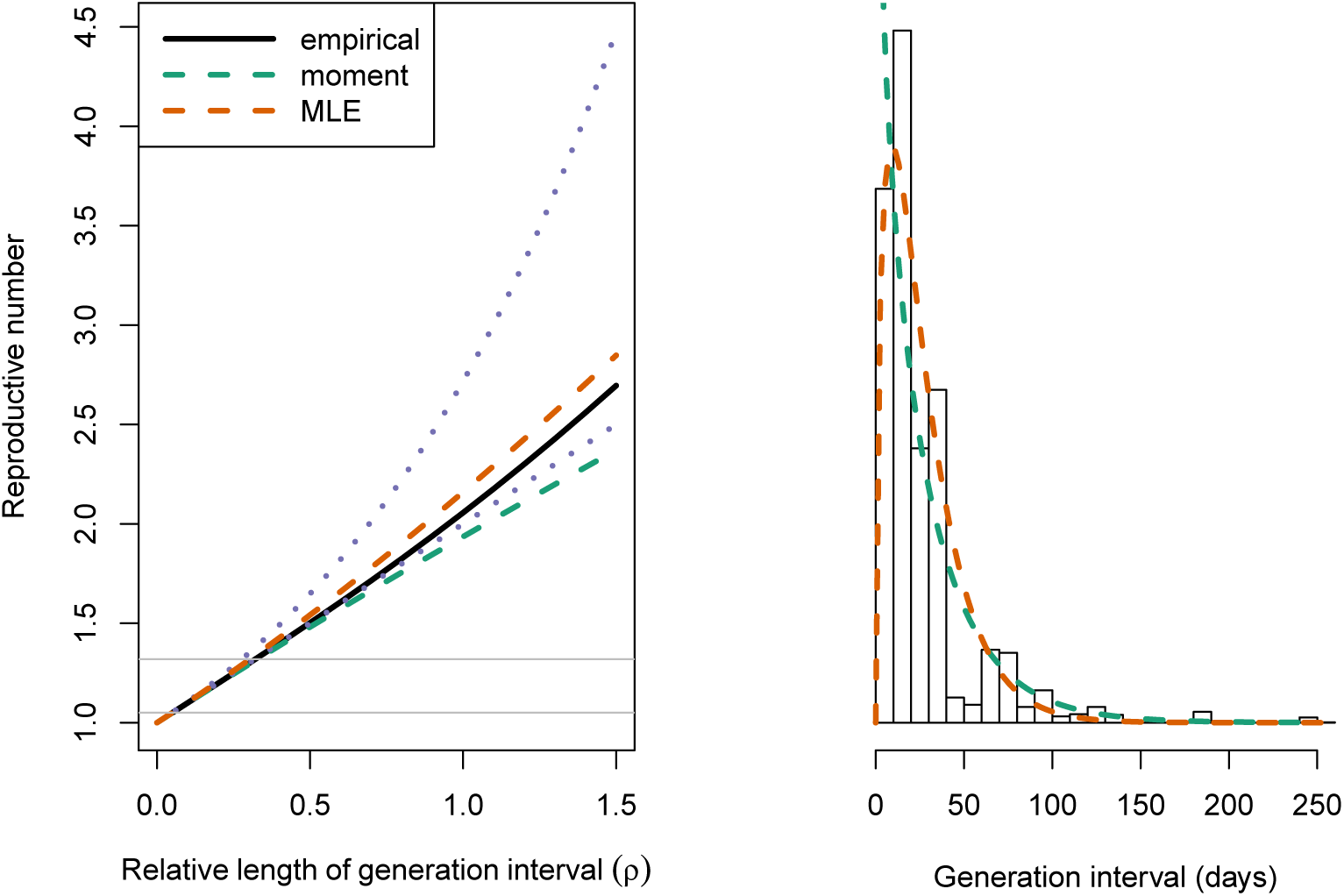
Estimating 𝓡 for the Rabies. (Left) estimating 𝓡 from rabies infectious case data. (solid curve) The exact speed-strength relationship (5) using a pseudo-realistic generation-interval distribution. (dashed curve) The Gamma approximated speed-strength relationship (6), using the mean and CV of a pseudo-realistic generation-interval distribution. “moment” approximation is based on the observed mean and CV of the distribution whereas “MLE” approximation uses the mean and CV calculated from a maximum-likelihood fit. (dotted curves) Naive approximations based on exponential (lower) and fixed (upper) generation distributions. Gray horizontal lines represent 𝓡 ranges estimated by [13]: 1.05-1.32. (Right) histogram represents rabies generation interval distributions simulated from incubation and infectious periods observed by [13]. Dashed curves represent estimated distribution of generation intervals using method of moments and MLE (corresponding to approximate speed-strength relationships of the left figure).

The reason for poor predictions of the moment approximation for higher 𝓡 can be seen in the histogram shown in Fig. 5. The moment approxima-tion is strongly influenced by rare, very long generation intervals, and does a poor job of matching the observed pattern of short generation intervals (in particular, the moment approximation misses the fact that the distribution has a density peak at a finite value). We expect short intervals to be par-ticularly important in driving the speed of the epidemic, and therefore in determining the relationship between *r* and 𝓡. We can address this problem by estimating gamma parameters formally using a maximum-likelihood fit to the pseudo-realistic generation intervals. This fit does a better job of match-ing the observed pattern of short generation intervals and of predicting the simulated relationship between *r* and 𝓡 across a broad range (Fig. 5).

## 5. Discussion

Estimating the reproductive number 𝓡 is a key part of characterizing and controlling infectious disease spread. The initial value of 𝓡 for an outbreak is often estimated by estimating the initial exponential rate of growth, and then using a generation-interval distribution to relate the two quantities [44, 39, 28, 40]. However, detailed estimates of the full generation interval are difficult to obtain, and the link between uncertainty in the generation interval and uncertainty in estimates of 𝓡 are often unclear. Here we introduced and analyzed a simple framework for *estimating* the relationship between 𝓡 and *r*, using only the estimated mean and CV of the generation interval. The framework is based on the gamma distribution. We used three disease examples to test the robustness of the framework. We also compared estimates based directly on estimated mean and variance of of the generation interval to estimates based on maximum-likelihood fits.

The gamma approximation for calculating 𝓡 from *r* was introduced by [29], and provides estimates that are simpler, more robust, and more realistic than those from normal approximations (see Appendix). Here, we presented the gamma approximation in a form conducive to intuitive understanding of the relationship between speed, *r*, and strength, 𝓡 (See Fig. 2). In doing so, we explained the general result that estimates of 𝓡 increase with mean generation time, but decrease with *variation* in generation times [44, 46, 37]. We also provided mechanistic interpretations: when generation intervals are longer, more infection is needed per generation (larger 𝓡) in order to produce a given rate of increase *r*. Similarly, when variance in generation time is large, there is more early infection. As early infections contribute most to growth of an epidemic, faster exponential growth is expected for a given value of 𝓡. Thus a higher value of 𝓡 will be needed to match a given value of *r*.

We tested the gamma approximation framework by applying it to parameter regimes based on three diseases: Ebola, measles, and rabies. We found that approximations based on observed moments closely match true answers (based on known, pseudo-realistic distributions, see Sec. 4 for details) when the generation-interval distribution is not too broad (as is the case for Ebola and measles, but not for rabies), but that using maximum likelihood to estimate the moments provides better estimates for a broader range of parameters Sec. 4.3, and also when data are limited (see Appendix).

Many studies have linked *r* with 𝓡 using generation interval distributions under various assumptions (Table 1). Simple methods, such as delta and exponential approximations, require minimal information (i.e., mean generation interval) but result in extreme estimates. More realistic methods, such as empirical and SEIR methods, may yield more realistic estimates, but are harder to interpret and require more data to implement.

Summarizing an entire generation interval distributions using two moments is a practically feasible approach that can give sensible and robust estimates of the relationship between *r* and 𝓡 that lie between extreme estimates (see Appendix). More detailed methods will often be preferred when data are abundant but the gamma approximation can be easily used in preliminary analyses. In particular, this framework has potential advantages for understanding the likely effects of parameter changes, and also for parameter estimation with uncertainty: since 𝓡 can be estimated from three simple quantities (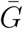, *κ* and *r*), it should be straightforward to propagate uncertainty from estimates of these quantities to estimates of 𝓡.

For example, during the Ebola outbreak in West Africa, many researchers tried to estimate 𝓡 from *r* [2, 7, 30, 35, 18] but uncertainty in the generation-interval distribution was often neglected (but see [41]). During the outbreak, used a generation-interval argument to show that neglecting the effects of post-burial transmission would be expected to lead to underestimates of 𝓡. Our generation interval framework provides a clear interpretation of this result: as long as post-burial transmission tends to increase generation intervals, it should result in higher estimates of 𝓡 for a given estimate of *r*. Knowing the exact shape of the generation interval distribution is difficult, but quantifying how various transmission routes and epidemic parameters affect the moments of the generation interval distribution will help researchers better understand and predict the scope of future outbreaks.

## Authors’ Contributions

SWP led the literature review, wrote the first draft of the MS, assisted with analytic calculations, and finalized the simulations and data analysis; JD conceived the study and did the initial simulations and analytic calculations;all authors contributed to refining study design, literature review, and MS writing. All authors gave final approval for publication.

## Competing interests

The authors declare that they have no competing interests.

## Acknowledgments

The authors thank Steve Bellan for helpful comments, and Katie Hampson for making rabies data available.

## Funding

This work was supported by the Canadian Institutes of Health Research [funding reference number 143486]. J.S.W. is supported by Army Research Office grant W911NF-14-1-0402.

## 6. Appendix

### 6.1 Comparison with the SEIR model

The SEIR (susceptible-exposed-infectious-recovered) model is given by

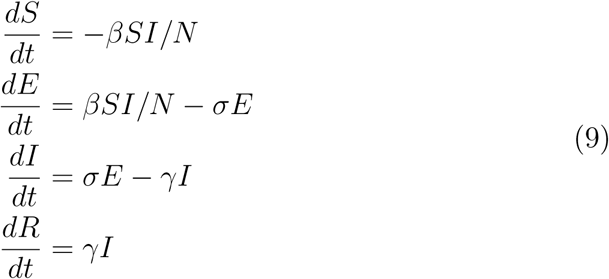

It is assume that latent and infectious periods are exponentially distributed with mean 1*/σ* and 1*/γ*, respectively.

For this model, generation interval distribution can be written as follows [39]:

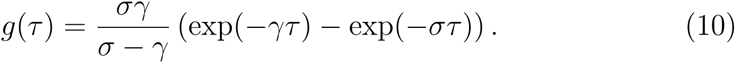

The mean generation interval is

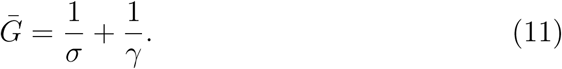

and the square coefficient of variation is

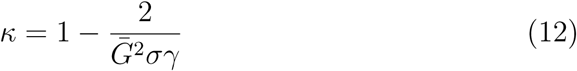

[21, 37] derived the following *r*–𝓡 relationship under SEIR model:

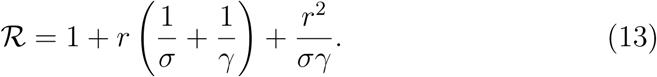

Reparameterizing, we have

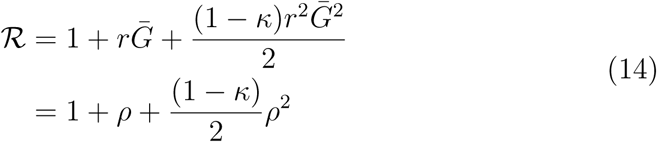

Note that Taylor expansion of (7) yields the following:

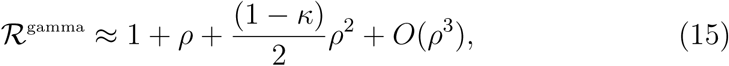

where 𝓡^gamma^ is an 𝓡 estimate based on the gamma approximation. When *σ* = *γ, g*(*τ*) follows a gamma distributio and *O*(*ρ*^3^) vanishes. We expect the gamma approximation to work well when average duration of latent and infectious periods are similar.

### 6.2 Ebola example

**Figure S1:**
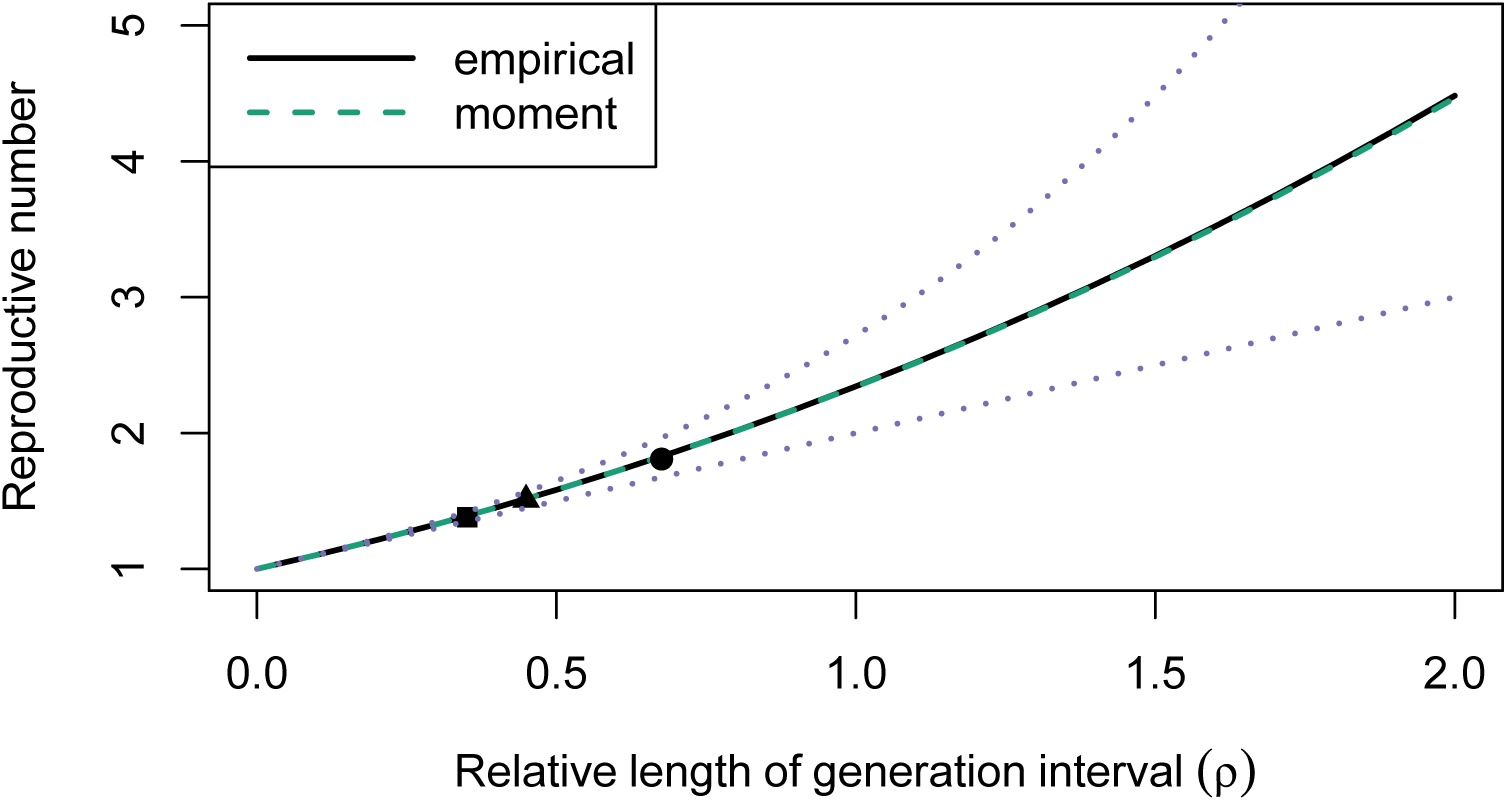
We perform the same analysis as we did in Sec. 4.1 assuming gamma distributed incubation and infectious periods. We find that the gamma approximated speed-strength relationship matches the true relationship almost perfectly in this case. Once again, we adjust the standard deviation of the incubation period to match the reported coefficient of variation in serial interval distributions. Rest of the parameters and points as in Fig. 3

### 6.3 Normal approximation

**Figure S2:**
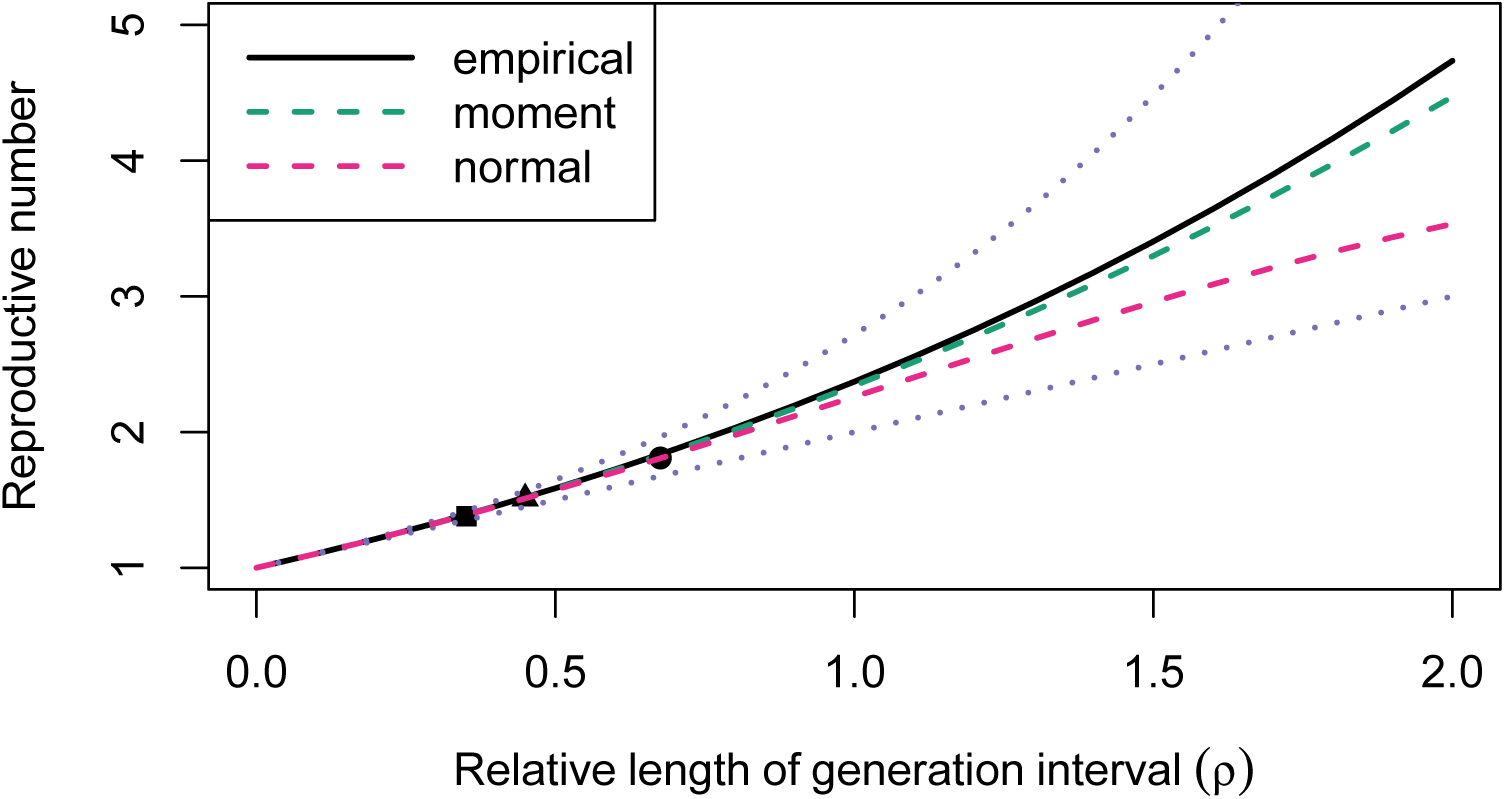
Approximating generation-interval distributions with a normal distribution has two problems. First, the distribution extends to negative values, which are biologically impossible. Second, as a consequence, the normal approximation predicts a saturating and eventually a decreasing *r–*𝓡 relationship for large *r*. Parameters and points as in Fig. 3.

### 6.4 Robustness of the gamma approximation

The moment-matching method (approximating 𝓡 based on estimated mean and variance of the generation interval) has an appealing simplicity, and works well for all of the actual disease parameters we tested (the breakdown for rabies distributions occurs for values of 𝓡 well above observed values). We therefore wanted to compare its robustness given small sample sizes along with that of the more sophisticated maximum likelihood method. Fig. S3 shows results of this experiment. When sample size is limited, estimates using MLE tend to be substantially close to the known true values in these experiments. As we increase sample size, our estimates become narrower. We also find that using the gamma approximated speed-strength relationship gives narrower estimates than the two naive estimates even when the sample size is extremely small (*n* = 10). It is important to note that Fig. S3 only conveys uncertainty in the estimate of coefficient of variation of generation interval distributions. Estimation of mean generation interval will introduce additional uncertainty into estimates of the reproductive number.

**Figure S3:**
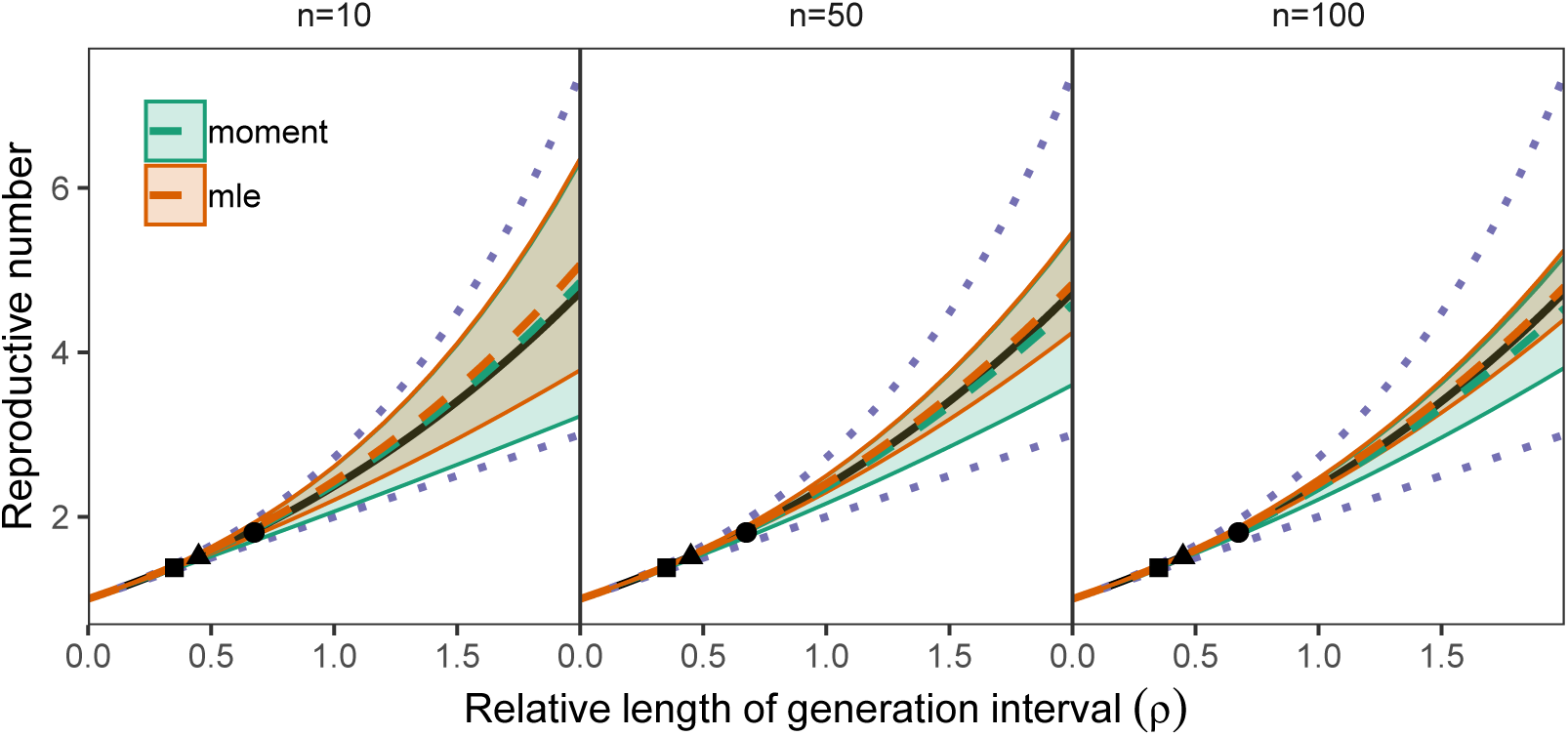
The effect of small sample size on approximated relationship between *r* and 𝓡. (black solid curve) The relationship between growth rate and 𝓡 using a known generation-interval distribution (see Fig. 3). (colored curves) Estimates based on finite samples from this distribution: dashed curves show the median and solid curves show 95% quantiles of 1000 sampling experiments. Note that the upper 95% quantile of the moment approximation and MLE approximation overlap. (dotted curves) Naive approximations based on exponential (lower) and fixed (upper) generation distributions.

